# Exquisite specificity of Pks13 defines the essentiality of trehalose in mycobacteria

**DOI:** 10.1101/2025.04.03.647003

**Authors:** Yushu Chen, Justin Dao Tian Lim, Shu-Sin Chng

## Abstract

Mycolic acids (MAs) are the major component of the mycobacterial outer membrane, a key contributor to the intrinsic resistance of mycobacteria to external insults including multiple antibiotics. After being synthesized in the cytoplasm and before reaching the outer membrane, MAs are transported across the inner membrane in the form of an acylated sugar, generally believed to be trehalose monomycolates (TMMs). Whether trehalose is the only mycolate carrier during transport is under debate, and why this highly abundant disaccharide is essential for mycobacterial growth is unclear. To address these questions, we leveraged a trehalose auxotrophic *Mycobacterium smegmatis* strain to investigate the biosynthetic steps affording TMMs. We show that in addition to TMMs, MAs are also not produced in the absence of trehalose. This is likely due to a product inhibition mechanism where un-reduced MA precursors accumulate on Pks13, the protein catalyzing the ligation of mycolic acids and the sugar head group. We establish that the un-reduced mycolates could only be released by trehalose, revealing exquisite Pks13 specificity, and subsequently reduced by CmrA in vitro. Furthermore, only trehalose and its analogs can reactivate MA biosynthesis in cells. Finally, by replacing trehalose with a 6-deoxy analog in cells, we demonstrate that the cord factor trehalose dimycolate is dispensable for *M. smegmatis* growth in vitro. Our work gives a clear depiction of how TMMs are formed and provides a compelling reason for the essentiality of trehalose, shedding light on the development of future anti-mycobacterial strategies.

**Significance Statement:** Tuberculosis and other diseases caused by mycobacteria pose a persistent threat to human health and well-being. The mycobacterial outer membrane is a highly impermeable barrier, conferring intrinsic resistance to multiple antibiotics. How the biosynthesis of mycolic acids, the major component of the mycobacterial outer membrane, is driven by trehalose is unknown. In this report, we study the molecular basis for the essentiality of trehalose. We define the order of two critical steps in the mycolic acid biosynthetic pathway involving trehalose and demonstrate the highly specific role that trehalose plays therein. This work answers key questions in terms of outer membrane biogenesis in mycobacteria, providing a venue for the development of novel anti-mycobacterial strategies.

## Introduction

Mycobacteria are the cause of a number of severe human diseases, such as tuberculosis (TB) and leprosy. There were more than 10 million new TB cases in a single year, and the disease alone claimed 1.1 million lives in 2023 (1). This situation is compounded by the spread of multi-drug resistant (MDR) *Mycobacterium tuberculosis* strains, and the higher risk of co-infection with TB in patients with human immunodeficiency virus. Non-tuberculous mycobacterial infections, caused by *Mycobacterium abscessus* and related organisms, have also been on the rise (2). The intrinsic resistance of mycobacteria to multiple antimicrobials and their emerging resistance to current drugs mark an urgent need for the identification of new drug targets.

Mycobacteria possess a distinct cell envelope. Besides the cytoplasmic membrane (IM) and the peptidoglycan (PG) cell wall, cells also contain an outer membrane (OM) bilayer, with the inner leaflet covalently linked to arabinogalactan (AG) polysaccharides extending outward from PGs (3, 4). The essential mycobacterial OM serves as an effective permeability barrier (5), which is largely contributed by the abundance of tightly packed mycolic acids (MAs) in the bilayer. These MAs are either linked to AG-PG as the entire inner leaflet (termed mycolyl-AGP or mAGP), or found as free lipids at the outer leaflet in the forms of trehalose monomycolates (TMMs) and trehalose dimycolates (TDMs). MAs are 2-alkyl, 3-hydroxyl, very long chain, branched fatty acids (C_58_-C_86_) present in mycobacteria, as well as phylogenetically related genera (6, 7). They comprise the shorter α branch (C_22_-C_24_) and the longer meromycolate chain (C_36_-C_60_), which are synthesized by a single FAS-I protein and a multi-protein FAS-II system, respectively (Fig. 1) (8). Different functional groups are introduced onto the meromycolate chain, giving rise to different MA classes - *M. tuberculosis* produces α-, methoxy-, and keto-MAs while *Mycobacterium smegmatis* contains α- (C_72_-C_84_) and epoxy-MAs, in addition to α’-MAs with a much shorter overall chain length (C_58_-C_66_) (8). Fully extended α-alkyl chains are carboxylated, and then condensed with meromycolate chains by polyketide synthase 13 (Pks13) (9), affording the MA precursors α-alkyl-β-ketoacids that are reduced by CmrA, (10) and esterified by trehalose to form TMMs (11); the order of these final two reactions in cells remain to be clarified (Fig. 1). TMMs are generally believed to be the final MA product in the IM and are continuously transported across the cell envelope to build the OM. The transport process is largely elusive, however, except for the identification of MmpL3 as the TMM flippase at the IM (12), and Ag85 enzymes as the mycolyltransferases for mAGP and TDM formation at the OM (13). Targeting the MA biosynthetic and transport pathways has proven to be an important anti-mycobacterial strategy, exemplified by the prevailing application of isoniazid, a first-line anti-TB drug targeting FAS-II (14).

**Figure 1.**
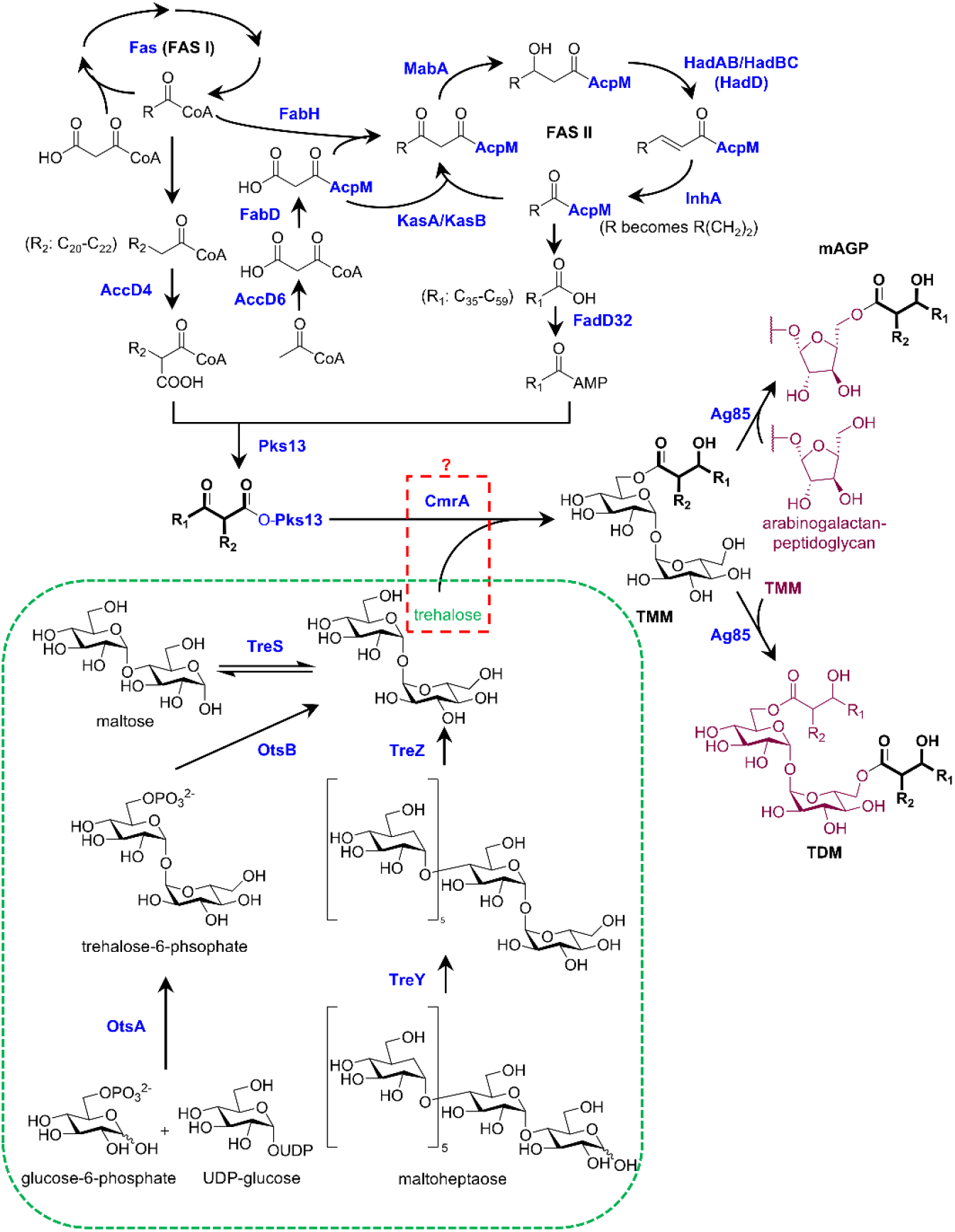
Biosynthetic pathways for trehalose, MAs, TMMs, TDMs and mAGP. Short chain α-branch fatty acids and long chain meromycolates are synthesized by FAS-I and FAS-II, respectively, and then condensed to give α-alkyl-β-ketomeromycolates on Pks 13. Subsequent trehalose-mediated release and CmrA-dependent reduction (in unknown order) of Pks13-linked α-alkyl-β-ketomeromycolates yields TMMs, which are transported to the OM and converted into TDMs and mAGP by Ag85 enzymes. FAS, fatty acid synthase.

There are knowledge gaps in mycolate biosynthesis that would be important for our understanding of mycobacterial physiology. One intriguing question is why trehalose is an essential metabolite for growth (15). It has been proposed that the role of trehalose as the final acceptor for MAs accounts for its essentiality. However, in vitro evidence suggested that Pks13 is not strictly specific for trehalose; several other mono- and disaccharides can release α-alkyl-β-ketoacids, albeit non-native short chain versions, from the enzyme to form the corresponding ester (11). Glycerol and glucose monomycolates have also previously been detected in lipidomic analyses (16, 17), although it is unclear whether they are derived directly from Pks13 or via mycolyl transfer reactions catalyzed by Ag85s, leading to sugar headgroup exchange (18). Therefore, is the predominance of TMM as the mycolate carrier in cells due to the high abundance of trehalose, or the hitherto unknown in vivo specificity of Pks13, which may then explain trehalose essentiality?

To elucidate acceptor specificity in cells, we need to devise a way to study the release of native mycolates from Pks13 by trehalose (or other sugars). We reasoned that we might be able to accumulate mycolates or its precursors on Pks13 by depleting the acceptor and stalling release. In this study, we exploited a trehalose auxotrophic *M. smegmatis* strain (TKO) lacking all three trehalose biosynthetic pathways (19), namely the OtsAB, TreYZ, and TreS pathways (20) (Fig. 1). This strain can only grow in the presence of exogenous trehalose actively taken up via the LpqY-SugABC ATP-binding-cassette (ABC) transporter (21). We conducted extensive lipid profiling of TKO cells depleted of trehalose, and reconstituted the final steps leading to TMM formation using native substrates from endogenously purified Pks13. We show that upon trehalose depletion, cells stop synthesizing mature MAs and TMMs, likely due to product accumulation on Pks13 that prevents further substrate turnover. Interestingly, only adding back trehalose, but not other sugars, can reactivate MA biosynthesis. Furthermore, we demonstrate that only trehalose can be used by purified Pks13 to release unreduced mycolates, for subsequent reduction by CmrA. Our work highlights the exquisite specificity of trehalose as the mycolate acceptor in cells, providing a molecular basis for its essentiality in mycobacteria.

## Results

### Trehalose depletion causes cessation of de novo MA biosynthesis

To study MA biosynthetic steps involving Pks13 and trehalose in mycobacteria, we first profiled newly synthesized lipids upon trehalose depletion in the *M. smegmatis* Δ*otsA*Δ*treYZ*Δ*treS* (TKO) trehalose auxotroph. TKO cells stopped growing after 48 h (*SI Appendix*, Fig. S1*A*), when we showed that intracellular trehalose was fully depleted (*SI Appendix*, Fig. S1*B*). At that point, cells were labeled with [^14^C]-acetate, and newly synthesized lipids in the forms of total extractable lipids (containing TMMs) or total mycolates (esterified as MA methyl esters, MAMEs) were isolated and analyzed by thin layered chromatography (TLC). As expected, mutant cells depleted of trehalose (dTKO cells) were deficient in TMM formation (Fig. 2*A*). Surprisingly, trehalose depletion also resulted in essentially no newly-synthesized mycolates (Fig. 2*B*), indicating that MA biosynthesis was possibly disrupted at steps before TMM formation. When the same cells were exogenously re-supplemented with trehalose during [^14^C]-labeling, both MA biosynthesis and TMM formation were restored (Fig. 2). Importantly, restoration of MA biosynthesis was not inhibited by several antibiotics that stopped cellular growth (*SI Appendix*, Fig. S2), suggesting that the cessation of MA synthesis was not due to growth arrest caused by lack of trehalose. The newly-synthesized MAs and TMMs, immediately upon trehalose re-supplementation, exhibited a higher α’ (C_58-66_) to α (C_72-84_) ratio than those in wild-type cells (Fig. 2*A* and *B*), a phenomenon also observed in cells when FAS-II was perturbed (*SI Appendix*, Fig. S3) (22-24). Transcriptomic analyses revealed that genes in the *fabD-acpM-kasA-kasB-accD6* operon, but not those encoding other FAS-II components, were up-regulated in trehalose-depleted cells (Fig. 3*A*), consistent with FAS-II perturbation. Furthermore, we observed up-regulation of the *acpS-fas, accD4-pks13-fadD32*, and *iniBAC* operons (Fig. 3*A* and *SI Appendix*, Fig. S4), perhaps a result of stalled MA biosynthesis, reminiscent of cellular responses of *M. tuberculosis* to FAS-I/II inhibitors (cerulenin, thiolactomycin and isoniazid) (25, 26). It is likely that TKO cells exhibit different metabolic states with or without trehalose. Following prolonged trehalose re-supplementation, the cellular transcriptomic profile of TKO cells became more similar to wild type cells (Fig. 3*A*). In addition, the ratio of α’-to α-mycolates was fully restored in a manner dependent on active transcription and translation (Fig. 3*B*–*D*). Taken together, our results establish that trehalose is required for de novo MA biosynthesis.

**Figure 2.**
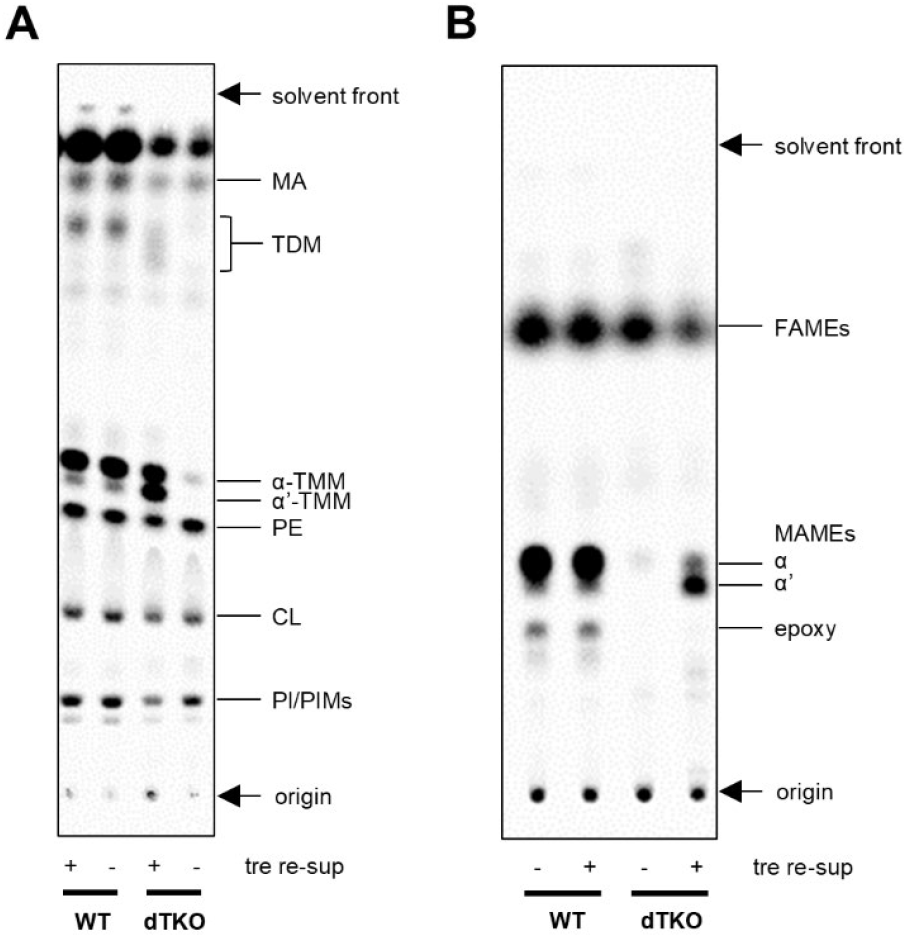
Trehalose is required for TMM formation and de novo MA biosynthesis. (*A*) TLC analysis of [^14^C]-labeled lipids extracted from *M. smegmatis* WT or dTKO (TKO depleted of trehalose) cells, with (+) or without (-) trehalose re-supplementation (tre re-sup). Developing solvent: chloroform:methanol:water 30:8:1. CL, cardiolipin; PE, phosphatidylethanolamine; PI, phosphatidylinositol; PIMs, phosphatidylinositol mannosides. (*B*) TLC analysis of [^14^C]-labeled fatty acids liberated from indicated strains, esterified as methyl esters. Three MAME species, α-, α’-, and epoxy-, were observed. Developing solvent: three times in hexane:ethyl acetate 95:5. TKO cells were depleted of trehalose (dTKO) prior to labelling and lipid extraction. FAME, fatty acid methyl ester; MAME, mycolic acid methyl ester. For samples on the same TLC plate, the same amount of radioactivity was loaded. The TLC plates were visualized by phosphor imaging.

**Figure 3.**
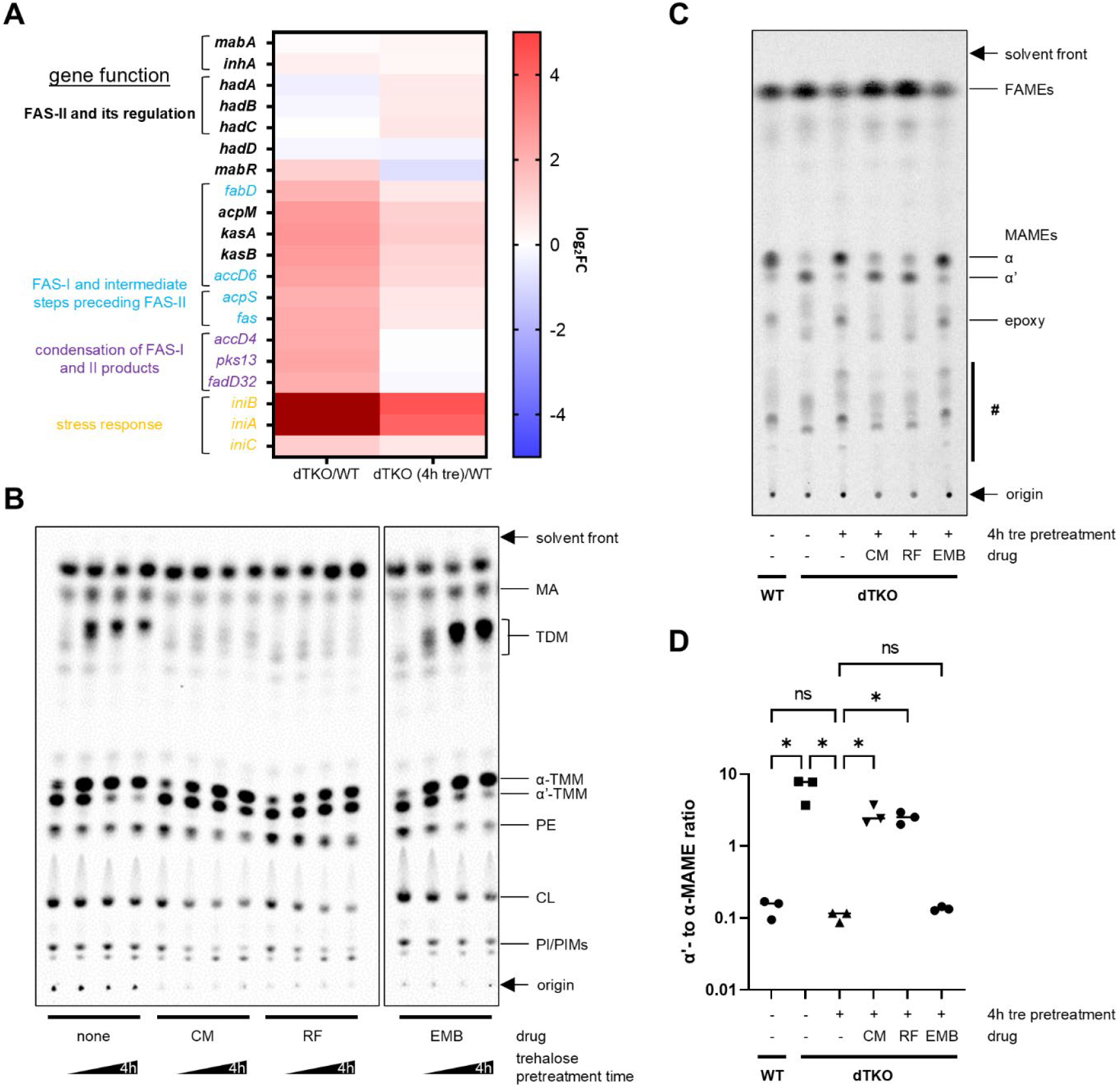
Trehalose-depleted TKO cells are in a distinct metabolic state. (*A*) Heat maps representing changes in transcript levels of indicated genes, either involved in MA metabolism or reportedly induced by FAS inhibitors, when comparing dTKO cells (trehalose-depleted TKO), or dTKO cells with trehalose re-supplementation (4h tre), to WT cells. Three biological replicates were used for each condition. FC, fold change. Maroon color on the heat map (*iniA* and *iniB* in the dTKO/WT column) indicates log_2_FC > 4. Genes in the same bracket are in the same operon. (*B*) TLC analysis of [^14^C]-labeled lipids extracted from dTKO cells pretreated with 20 μM trehalose in the presence of different drugs for extended periods of time (0, 1, 2.5 or 4 h) before radiolabeling with [^14^C]-acetate. Developing solvent: chloroform:methanol:water 30:8:1. Chloramphenicol (CM), rifampicin (RF) and ethambutol (EMB) inhibit translation, transcription and AG biosynthesis, respectively. CL, cardiolipin; PE, phosphatidylethanolamine; PI, phosphatidylinositol; PIMs, phosphatidylinositol mannosides. (*C*) TLC analysis of [^14^C]-labeled total acyl methyl esters derived from dTKO cells pretreated with 20 μM trehalose in the presence of different drugs for 4 h before radiolabeling with [^14^C]-acetate. “#” denotes lipids with unknown identities. Developing solvent: three times in hexane:ethyl acetate 95:5. For samples on the same TLC plate, the same amount of radioactivity was loaded. The TLC plates were visualized by phosphor imaging. (*D*) Quantification of α’- and α-MAME in (*C*). Each data point (mean ± SD) represents the results from three biological replicates. ns, not significant; *, p < 0.05. FAME, fatty acid methyl ester; MAME, mycolic acid methyl ester.

### Trehalose depletion causes accumulation of keto-mycolates on Pks13 that must be released and reduced sequentially to form TMMs

Both mycolate biosynthesis and TMM formation were abolished in trehalose-depleted cells. We reasoned that these observations may indicate accumulation of lipids on Pks13, which prevents further turnover of mycolate substrates. To test this idea, we labeled *M. smegmatis* dTKO cells with [^14^C]-acetate and analyzed the whole cell protein profile via SDS-PAGE. Notably, a radioactive band showed up at ∼200 kDa (*SI Appendix*, Fig. S5), which is consistent with the size of Pks13 (197.1 kDa). The radioactivity on the band could be reduced by treatment with either trehalose or the mycolate biosynthesis inhibitor isoniazid (*SI Appendix*, Fig. S5), suggesting that the labeled band contains mycolate-related lipids. We next expressed C-terminally His-tagged Pks13 from an integrative plasmid, and isolated it from [^14^C]-acetate-labeled dTKO cells via affinity purification. Remarkably, we detected significant radioactivity on Pks13-His from these cells (Fig. 4*A*); the signal was much weaker when the same cells had been treated with trehalose or isoniazid (Fig. 4*B* and *C*). These results indicate that mycolate precursors accumulate on Pks13 upon trehalose depletion, likely accounting for why de novo biosynthesis of mycolates is abolished in the absence of trehalose.

**Figure 4.**
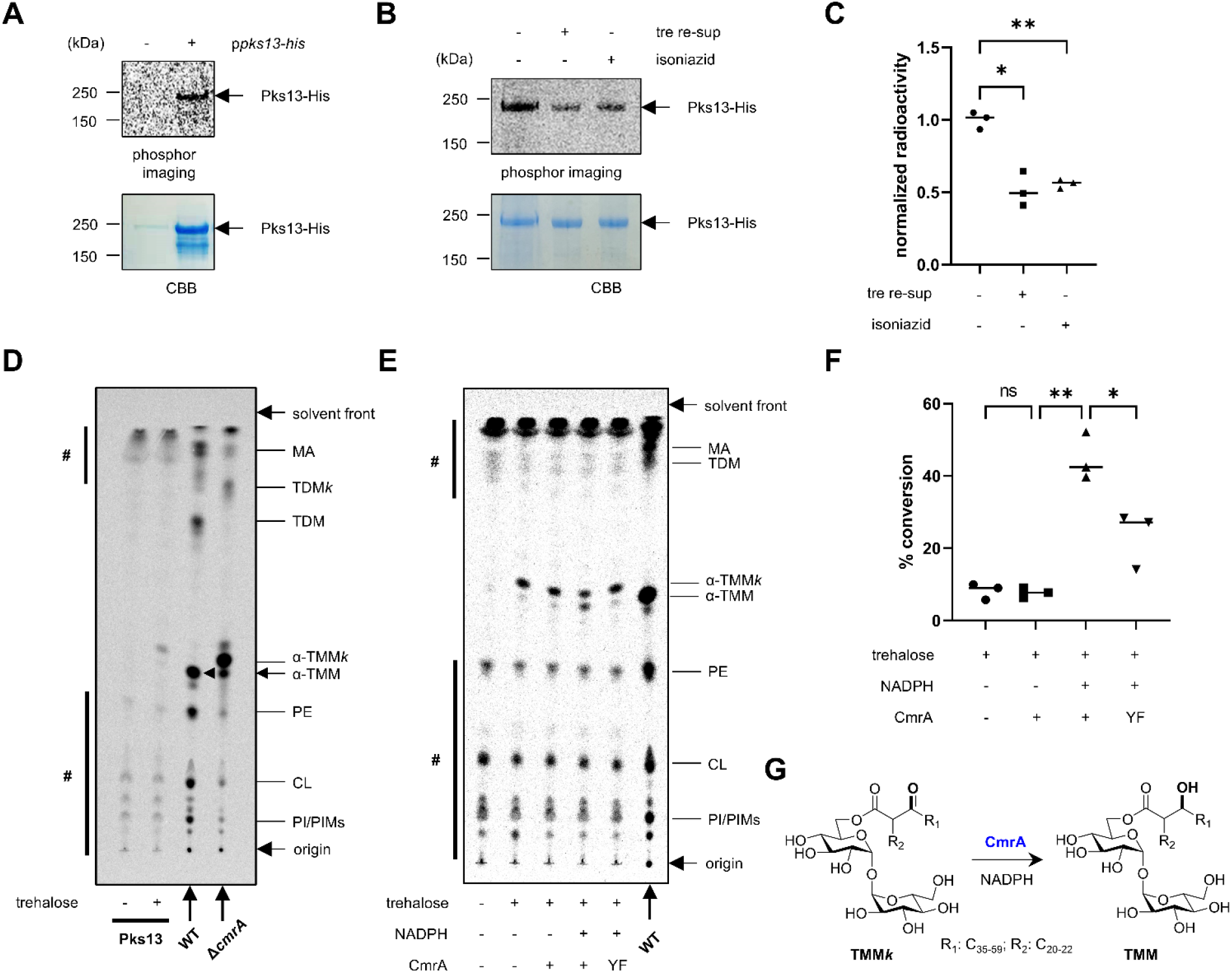
Unreduced mycolates accumulate on Pks13 upon trehalose depletion. *(A)* and *(B)* SDS-PAGE analyses of Pks13-His affinity purified from dTKO cells harboring empty vector or pJEB402-*pks13*-*his*_*8*_ grown in the presence of [^14^C]-acetate. The gel was visualized by phosphor imaging (top) and Coomassie brilliant blue staining (CBB; bottom). tre re-sup, trehalose re-supplementation. In *(B)*, trehalose (tre re-sup) or isoniazid was added together with [^14^C]-acetate. (*C*) Quantification of the Pks13-associated lipids in different conditions in (*B*). Each data point (mean ± SD) represents the results from three biological replicates. *, p < 0.05; **, p < 0.01. (*D*) and (*E*) TLC analyses of [^14^C]-labeled lipids extracted from affinity-purified Pks13-His subjected to (*D*) in vitro mycolate release by trehalose, and (*E*) reduction by CmrA. Lipids extracted from WT and Δ*cmrA* cells were used as references for migration positions of α-TMM and α-TMM*k*, respectively. For clarity, α’-TMM and α’-TMM*k* (spots immediately below corresponding α-TMM and α-TMM*k*) are not labelled for these samples. CL, cardiolipin; PE, phosphatidylethanolamine; PI, phosphatidylinositol; PIMs, phosphatidylinositol mannosides. For in vitro samples, the top and bottom spots (#) on the TLC other than TMM*k* and TMM are present even in the absence of trehalose and are likely non-specific lipids bound to TALON resin used for affinity purification of the protein. Developing solvent: chloroform:methanol:water 30:8:1. The TLC plates were visualized by phosphor imaging. (*F*) Quantification of the conversion of trehalose mono-β-ketomycolate (TMM*k*) to TMM in different conditions in (*E*). Each data point (mean ± SD) represents the results from three biological replicates (see *SI Appendix*, Fig. S6 for other replicates). YF, CmrA containing a Y157F mutation. ns, not significant; *, p < 0.05; **, p < 0.01. (*G*) Reaction scheme illustrating CmrA-catalyzed reduction of TMM*k* affording TMM.

To elucidate what lipid species may have accumulated on Pks13, we attempted to use trehalose to release lipids from affinity-purified Pks13-His in vitro. Interestingly, trehalose indeed facilitated the formation of a new extractable lipid species in the reaction mixture. The lipid migrated slightly faster than TMM on normal-phase TLC, suggesting that it is a less polar derivative of the latter (Fig. 4*D*). By comparing the mobility of this lipid to TMM species synthesized in a strain lacking the reductase CmrA (10), we assigned it to be trehalose α-alkyl-β-ketomeromycolate, the unreduced keto form of TMM (TMM*k*). The release of the α-alkyl-β-ketomeromycolate precursor to form TMM*k* is believed to be catalyzed by the thioesterase domain of Pks13. While this reaction can happen in the absence of the ketoreductase CmrA both in vitro (11) and in cells (27), it is not clear whether the reduction step actually takes place before or after the release. Notably, the observed accumulation of α-alkyl-β-ketomeromycolates in the absence of trehalose demonstrates the inability of endogenous CmrA to reduce this precursor directly on Pks13. Consistent with this idea, only when we also added purified CmrA and NADPH during the reaction, we observed the conversion of TMM*k* to mature TMM (Fig. 4*E*–*G* and *SI Appendix*, Fig. S6). A mutation at the putative CmrA active site (CmrA^Y157F^) reduced the conversion significantly. Therefore, we have effectively recapitulated in vitro the final release and reduction steps in the formation of TMM. We conclude that trehalose release of mycolates from Pks13 allows subsequent CmrA reduction to form TMMs.

### Release of native ketomycolates from Pks13 is highly specific to trehalose

Having established that unreduced MA precursors can be released from Pks13 and esterified by trehalose in vitro, we further explored if other sugars could act as MA acceptors. In an earlier study, unreduced MA analogs (C_30_) could be synthesized in vitro and released from recombinant Pks13 by a few mono- and disaccharides other than trehalose, including mannose, allose, maltose, and most prominently glucose (11) (Fig. 5*A*). Surprisingly, none of these sugars facilitated formation of ketomycolate sugar esters when added to Pks13 endogenously purified from dTKO cells (Fig. 5*B*); we reasoned that the longer chain length of native ketomycolates can in fact contribute to the sugar specificity of the MA acceptor. We also tested other trehalose analogs in the release assay. Notably, 6-azido-trehalose, but not 6,6’-diazido-trehalose, could drive the formation of 6-azido-TMM*k* in the reaction mixture, consistent with the requirement of a free 6’-OH group; this observation also corroborates that α-alkyl-β-ketomeromycolates accumulated on Pks13 in the absence of trehalose. The identity of the 6-azido-trehalose-released product was assigned based on its similar migration on TLC when compared to 6-azido-TMM*k* produced in Δ*cmrA* cells fed with 6-azido-trehalose (*SI Appendix*, Fig. S7). To extend the finding on acceptor specificity, we next examined the ability of the various sugars in activating de novo mycolate biosynthesis when re-supplemented to dTKO cells. Remarkably, only 6-azido-trehalose (in addition to trehalose) enabled the formation of newly-synthesized MAs (Fig. 5*C*), and thus TMMs (*SI Appendix*, Fig. S8*A*), consistent with its ability to release accumulated precursors from Pks13. We confirmed the assignment of 6-azido-TMMs hence formed by demonstrating its reactivity with an alkyne-containing probe via click chemistry (*SI Appendix*, Fig. S8*B*) (28). We conclude that Pks13 exclusively uses trehalose as the MA acceptor in cells, which can explain the essentiality of trehalose in mycobacteria.

**Figure 5.**
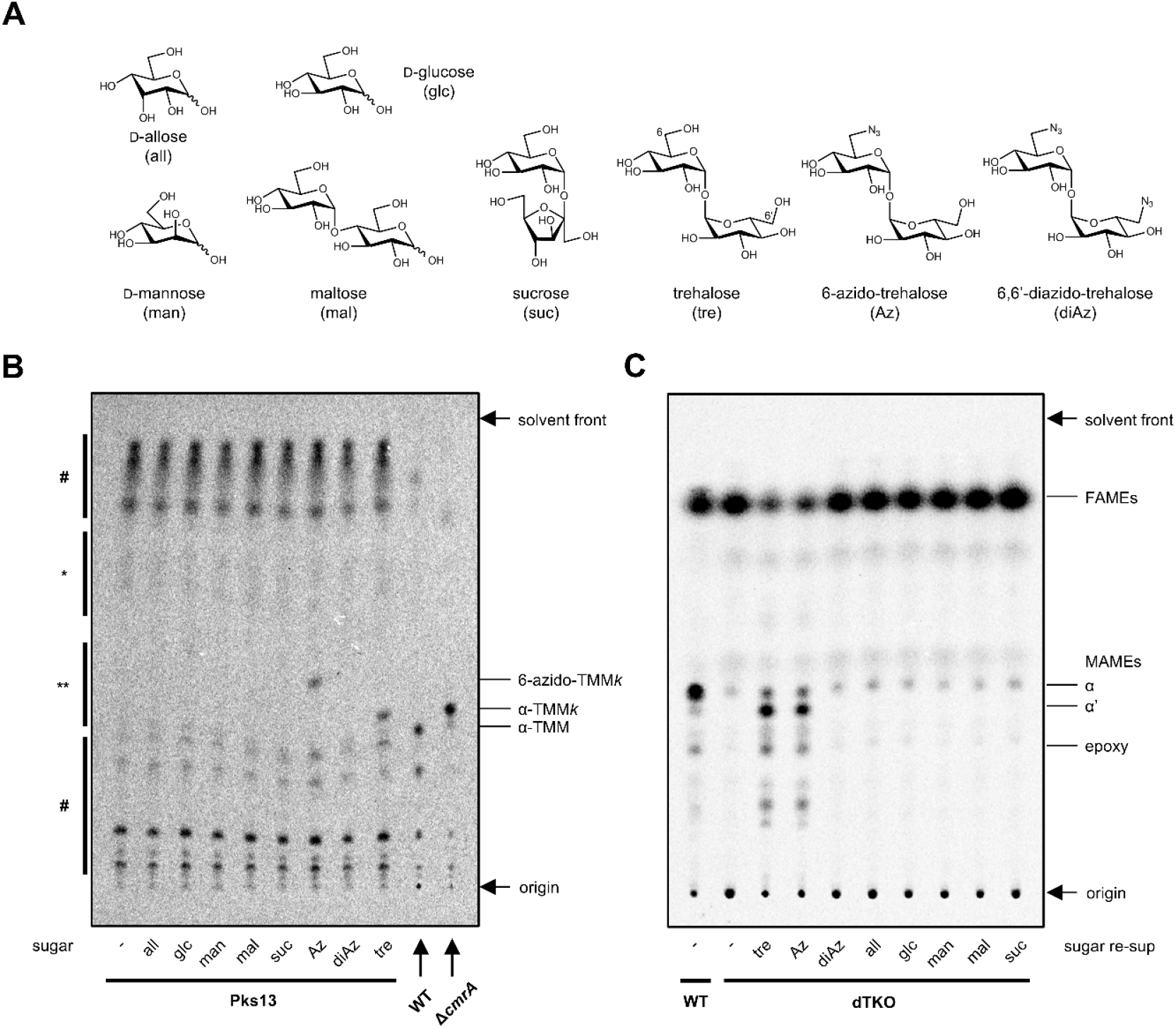
Pks13 specifically uses trehalose to esterify MAs. (*A*) Chemical structures of sugars tested. (*B*) TLC analysis of lipids extracted from the same amount of affinity-purified Pks13-His incubated with the indicated sugar. Total extractable lipids from WT and Δ*cmrA* cells were used as references for migration positions of TMM and TMM*k*, respectively. The top and bottom spots (#) on the TLC are likely non-specifically bound to TALON resin used for affinity purification of the protein. *, a region where monosaccharide monomycolates are expected to appear; **, a region where disaccharide monomycolates are expected to appear. (*C*) TLC analysis of [^14^C]-labeled total acyl methyl esters derived from dTKO cells re-supplemented with the indicated sugar. Developing solvent: three times in hexane:ethyl acetate 95:5. The TLC plates were visualized by phosphor imaging.

### TDM is dispensable for *M. smegmatis* growth in vitro

It is intriguing that 6-azido-trehalose is able to release mycolate precursors from Pks13 (Fig. 5*B*) and facilitate continuous MA biosynthesis (Fig. 5*C*). In fact, we found 6-azido-trehalose alone could support the growth of TKO cells, albeit at a slower rate than that supported by the same amount of trehalose (Fig. 6*A*). This observation implies that *M. smegmatis* cells can survive without TDM; in the trehalose auxotrophic strain grown in 6-azido-trehalose, mAGP but not TDM could be formed, since the mycolyl chain from 6-azido-TMM can still be transferred to AGs but not another 6-azido-TMM molecule lacking the 6-OH group (Fig. 1). Using [^14^C]-acetate lipid profiling, we confirmed that 6-azido-trehalose-fed TKO cells made mAGP but not TDM (Fig. 6*B*). Notably, we did not detect any major difference in drug sensitivity between TKO cells with and without TDM (*SI Appendix*, Table S1). Therefore, we assert that the TDM, despite its key role in pathogenicity as the cording factor (29), is not essential for growth and OM permeability function of *M. smegmatis* in vitro, and possibly other mycobacteria.

**Figure 6.**
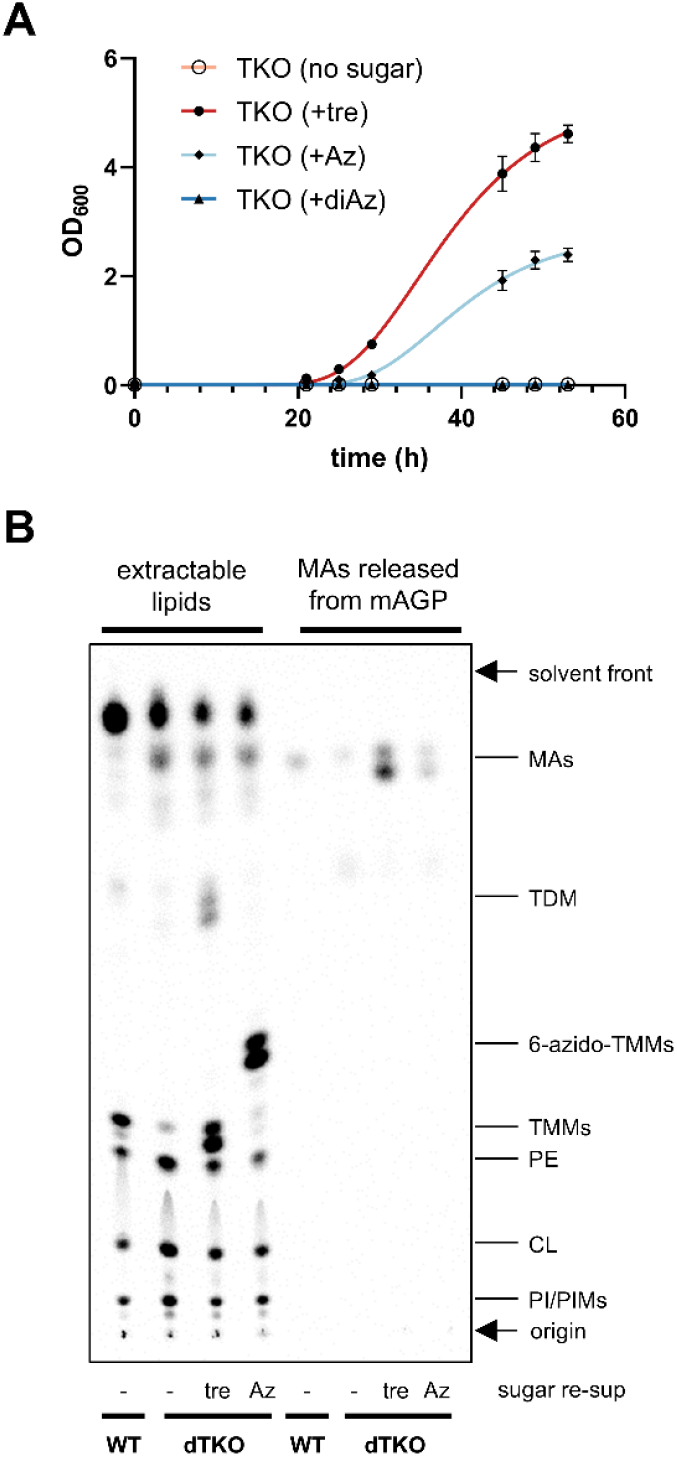
TDM is not required for mycobacterial growth in vitro. (*A*) Growth curves of TKO cells in the presence of the indicated trehalose analogs. TKO cells were depleted of trehalose before subculture into fresh medium (no sugar) or that containing 5 μM trehalose (+tre), 6-azido-trehalose (+Az) or 6,6’-diazido-trehalose (+diAz). Each data point (mean ± SD) represents the results from three biological replicates. (*B*) TLC analysis of [^14^C]-labeled lipids extracted from trehalose-depleted (dTKO) cells re-supplemented with trehalose (tre) or 6-azido-trehalose (Az). TMMs and TDMs are fully extracted from cells, while MAs conjugated as mAGP were released by base hydrolysis. Developing solvent: chloroform:methanol:water 30:8:1. The same amount of radioactivity was loaded for each extractable lipid sample, while MAs released from mAGP were loaded in the same volume as extractable lipids from the same cells. The TLC plate was visualized by phosphor imaging.

## Discussion

By using a trehalose auxotrophic strain, we have demonstrated that trehalose depletion abolishes de novo MA biosynthesis, likely due to intermediates accumulating on Pks13. In addition, we have established that Pks13 is highly specific, and only uses trehalose to release β-keto-mycolates from its thioesterase domain. Furthermore, we have found that CmrA is unable to reduce β-keto-mycolates on Pks13 itself in cells, a process that occurs only after the formation of TMM*k*. Our study has recapitulated the final steps in the MA biosynthetic pathway with native substrates, and the details therein, revealing why the specific disaccharide trehalose is essential in mycobacteria.

Our findings that trehalose is a highly specific acceptor for mycolates on Pks13 differ from an earlier study in vitro (11). One key factor that may account for this difference is the use of short mycolate analogs in that study, compared to the tracking of native substrates accumulating on Pks13 here, both in cells and in vitro. The thioesterase domain in polyketide synthases (PKSs) often serves as a chain length regulator or a proof-reading enzyme: in the case of shorter products formed on the acyl carrier protein domain, the thioesterase domain may remove the acyl chain to prevent blockage of the synthetic cascade (30). We therefore reason that the use of shorter mycolate analogs may have rendered Pks13 more promiscuous in the choice of the sugar acceptor. Furthermore, it is conceivable that high specificity for trehalose as the acceptor for mycolates is important for streamlining downstream processes, at least for transport. Specifically, the exquisite synthesis of TMM circumvents the need for multiple redundant transporters to accommodate different hydrophilic sugar headgroups during flipping across the IM, and/or transport to the OM (31).

Mycobacteria leverage multiple PKSs for the biosynthesis of complex lipids to build their unique OM (32). Among them, Pks13 is essential for mycobacterial growth (9) and has been revealed as the target of a few hit and lead compounds (33-35). It had not been established whether the reduction of ketomycolates by CmrA happens before or after the release and esterification by trehalose in the TMM biosynthesis pathway. Cells lacking CmrA still produce TMM*k* (10), indicating that ketomycolates can be released from Pks13 by endogenous trehalose. Notably, we have now demonstrated that cells depleted of trehalose accumulate ketomycolates on Pks13, revealing that endogenous CmrA cannot reduce this precursor. We have further shown that CmrA-mediated reduction would proceed after the formation of TMM*k* in vitro, establishing that reduction of ketomycolates only takes place after release from Pks13 by trehalose. Such post-release reduction contrasts from most other PKSs where keto-reduction steps occur before final release (36). Given that β-keto-acids can undergo spontaneous decarboxylation reactions, we speculate that preventing reduction of ketomycolates in the absence of trehalose may provide an avenue for degradation and re-assimilation, especially in the context of corynebacteria where MAs are non-essential (37).

We have shown that trehalose is required for de novo MA biosynthesis. A simple explanation is that the accumulation of intermediates on Pks13 without trehalose prevents reduction by CmrA, and thus formation of mature mycolates. However, the impact of such accumulation on substrate loading and condensation on Pks13, or on upstream MA biosynthetic steps, cannot be ruled out. It is also not clear how inhibition of MA biosynthesis upon trehalose depletion results in a transcriptional response that up-regulates *acpS-fas, fabD-acpM-kasA-kasB-accD6* and *accD4-pks13-fadD32* operons, which likely accounts for the high α’-to α-MA ratio when trehalose was re-supplemented. Interestingly, these transcriptomic changes are highly similar to those reported for drugs inhibiting FAS-I or II enzymes that are responsible for mycolic acid biosynthesis, but not other envelope pathways (25), suggesting that the changes were elicited specifically by lack of mature mycolates. While it is clear that cells depleted of trehalose exhibit general envelope stress, evidenced by up-regulation of the *iniBAC* operon (26), we postulate that mycobacteria may have yet-to-be-identified mechanisms to sense mycolates in the cell, so as to counteract defects in overall envelope assembly. This would be analogous to how Gram-negative bacteria deploy various LPS sensing mechanisms to ensure proper OM assembly (38).

Finally, we have also been able to show that TDM is not essential for in vitro mycobacterial growth. While other studies have alluded to this idea, it was never before possible to remove TDM completely (39, 40); this limitation may also impact the interpretation of in vivo studies that ascribed TDM as a key virulence factor and immune response modulator (41, 42). A trehalose auxotrophic strain supplemented with 6-deoxy-trehalose analogs may thus be useful for future studies on TDM in vivo. Our study provides valuable insights into the formation of MA, TMM and TDM, and serves as a foundation for the development of new anti-mycobacterial drugs, especially against Pks13.

## Supporting Information

- Detailed materials and methods.
- Figures S1-S8 and Table S1

## Acknowledgments

We thank Prof. Carolyn Bertozzi (Stanford University) and Prof. Rainer Kalscheuer (Heinrich-Heine-Universität Düsseldorf) for providing the TKO strain, as well as various single and double mutants in trehalose biosynthesis. 6-azido-trehalose is a kind gift from Prof. Benjamin Swarts (Central Michigan University). We thank Dr. Qiping Liu (Chemical, Molecular and Materials Analysis Centre, National University of Singapore) for the guidance and help with GC-MS. Library preparation and RNA sequencing was done by the integrated Genome Analytics Platform at the Genome Institute Singapore. This project was funded by Singapore Ministry of Education Academic Research Fund Tier 2 grant MOE-000116 (S.-S.C.).

## Notes

### Competing Interest Statement

The authors have declared no competing interest.

### Summary of Updates

Figures 1, 3, 4, and 5 updated; new Figures S3, S7, S8; more text amendments

