## Supporting Information for "Exquisite specificity of Pks13 defines the essentiality of trehalose in mycobacteria"

**This PDF file includes**

Materials and Methods

Table S1

Figures S1 to S8

### Materials and Methods

#### Bacterial strains and growth conditions

The TKO strain was derived from the *M. smegmatis* mc$^2$155 wild type strain and was obtained from Prof. Rainer Kalscheuer (1). *M. smegmatis* cells were grown in 7H9 medium (BD Biosciences) containing 0.5% glycerol and 0.05% tyloxapol, except for the Δ*cmrA* strain, which was grown in tryptic soy broth (BD Biosciences) containing 0.05% Tween 80. To allow depletion of trehalose from the cells, 1 µM trehalose (Alfa Aesar) was added into inoculated TKO liquid culture and grown for 48 h. The cells collected at this point were referred to as dTKO (trehalose-depleted TKO) cells. The *E. coli* NovaBlue strain was used for cloning. *E. coli* cells were grown in LB-Miller medium (BD Biosciences). LB plates containing 1.5% agar (BD Biosciences) were used as solid media for plasmid construction and cell growth for both *E. coli* and *M. smegmatis*. Where needed, 50 µg/mL hygromycin (MilliporeSigma) and 20 µg/mL kanamycin (MilliporeSigma) were used for *M. smegmatis*. Hygromycin, kanamycin, spectinomycin (MilliporeSigma), and ampicillin (MilliporeSigma) were used at 150 µg/mL, 25 µg/mL, 30 µg/mL, and 100 µg/mL respectively for *E. coli*.

The Δ*cmrA* strain was constructed by two-step homologous recombination (2). Briefly, a suicide plasmid pYUB854 with no replication origin and a hygR cassette was used as the backbone for the insertion of two flanking regions of the target gene side by side, and a *lacZ-sacB* cassette for blue-white screening and negative selection on sucrose. The plasmid was electroporated at 2250 V using Eporator (Eppendorf) into *M. smegmatis* mc$^2$155 electrocompetent cells, which were subsequently plated on LB plates containing 50 µg/mL hygromycin (Merck, 400051) and 20 µg/mL 5-bromo-4-chloro-3-indolyl-β-D-galactopyranoside (X-gal; 1st BASE, BIO-1020). The plate was incubated at 37 ºC for 5 days to allow the growth of the first cross-over cells (those forming blue colonies on the plate). Cells harboring the first cross-over at the targeted site were grown in Middlebrook 7H9 broth (Becton Dickinson, 271310) containing 0.5% glycerol and 0.05% tyloxapol (Merck, T0307) and spread onto LB plates containing 20 µg/mL X-gal and 5% sucrose. The plate was again incubated at 37 ºC for 4 days to allow the growth of second cross-overs (those forming whitish-yellow colonies on the plate). Correct gene deletion mutants were screened by PCR and Sanger sequencing (Bio Basic Asia Pacific) among the second cross-overs.

**Quantification of intracellular trehalose**

The quantification of intracellular trehalose was done following methods in (3) with slight modification. *M. smegmatis* TKO cells depleted of trehalose (dTKO) were harvested after 48 h growth by centrifugation at 4,000 x *g* for 5 min. The amount of cells harvested was equivalent to that in a 10-mL culture where $OD_{600}$ = 1.0. After the supernatant was removed, the cells were washed once with 1x PBS pH 7.4. The pellet was incubated in 1 mL 50% ethanol (v/v) in water at 80 °C for 30 min to lyse the cells. The solution was then dried using a SpeedVac concentrator. The sample was incubated in 500 µL acetic anhydride and 500 µL pyridine at 95 °C for 20 min to acetylate free trehalose. As an external standard, a specific amount of trehalose (Alfa Aesar) was acetylated using the same procedure. Acetylated trehalose was dried and dissolved in 200 µL acetone for GC-MS analysis. The analysis was performed on a Shimazu QP2010 Plus GC-MS system equipped with an HP-5MS (30 m x 250 µm x 0.25 µm) column. 2 µL sample was injected with a 1:1 split ratio. The column temperature was programmed as follows: 50 °C for 1 min, 30 °C/min to 200 °C, then 8 °C/min to 320 °C and 320 °C for 8 min. The interface temperature between GC and MS was 280 °C. The mass spectrometer was set to scan from 50 to 500 amu. The quantification was done by comparing the peak area given by GC with that of the acetylated trehalose standard.

**Lipid extraction and TLC analysis**

*M. smegmatis* TKO cells were grown to stationary phase to deplete trehalose (dTKO). 1 mL of the culture was pelleted at 4,000 x *g* for 3 min, resuspended in fresh 7H9 medium, and incubated with 0.2 µCi/mL [1-$^{14}$C]-acetate (Perkin Elmer NEC084A001MC) for 45 min to label the cellular lipids. Different sugars were used at 20 µM. The labeled cells were harvested by centrifugation at 4,000 x *g* for 3 min.

For analysis of extractable lipids, the cells were lysed and extracted in 750 µL chloroform:methanol

(2:1, v/v) by sonication in a water bath. The sonication was performed three times, each 30 s with 5 s vortexing in between. 250 μL methanol and 350 μL ultrapure water were then added to the mixture, which was subjected to centrifugation at 10,000 x *g* for 3 min to allow phase separation. The bottom organic phase was collected and dried in a fumehood overnight. For DBCO-acid treated samples, the sample was resuspended in 100 μL chloroform, incubated with 0.3 mM DBCO-acid (MilliporeSigma) for 1 h at room temperature and dried again. The lipids were dissolved in 50 μL chloroform:methanol (2:1, v/v) for TLC analysis. The total radioactivity of each sample (cpm) was measured using a MicroBeta2 scintillation counter. Lipids of the same amount of radioactivity (20,000 cpm) were spotted onto silica gel 60 $F_{254}$ glass TLC plates (MilliporeSigma). The plate was developed once with a chloroform:methanol:water (30:8:1, v/v/v) solvent system in a glass chamber pre-equilibrated for 30 min with the solvent. After development, the TLC plate was completely dried in a fumehood for 1 h. It was then exposed to a storage phosphor screen for 3 days followed by visualization by phosphor imaging (Storm, GE Healthcare).

For analysis of mycolate released from mAGP, the top aqueous phase after lipid extraction was extracted again with 500 μL chloroform to remove residual extractable lipids. The aqueous phase was then dried in a speed-vac Concentrator Plus (Eppendorf). MAs were liberated from the dried cell debris by heating at 90 ˚C in 13% (w/v) tetrabutylammonium hydroxide (TBAH; MilliporeSigma) for 2 hours. After cooling down to room temperature, it was acidified to pH around 4 with concentrated hydrochloric acid. Free MAs were extracted twice with 500 μL *n*-hexane each time. The pooled sample was dried in a fumehood overnight, dissolved in chloroform:methanol (2:1, v/v) for TLC analysis. The plate was developed once with a chloroform:methanol:water (30:8:1, v/v/v) solvent system.

For total fatty acid and mycolic acid methyl esters (FAMEs and MAMEs), the cells were resuspended in 400 μL 13% (w/v) TBAH before the whole mixture was heated at 90 °C for 2 h. After cooling, 200 μL ultrapure water, 400 μL dichloromethane, and 30 μL iodomethane (MilliporeSigma) were sequentially added. The mixture was left on an orbital shaker for 1 h to allow methyl ester formation. The resulting mixture was separated into two phases by centrifugation at 10,000 x *g* for 2 min. The bottom organic layer was collected and dried in a fumehood overnight. The MAMEs were dissolved in 50 μL chloroform analyzed by TLC. Lipids of the same amount of radioactivity (20,000 cpm) were spotted onto the plate, which plate was developed thrice in a hexane:ethyl acetate (95:5, v/v) solvent system.

**RNA extraction**

One volume of RNAprotect Bacteria Reagent (Qiagen) was added to mycobacterial liquid cultures of total $OD_{600}$ 15. The mixture was incubated for 5 min at room temperature after brief vortexing. The cells were harvested by centrifugation at 4,000 x *g* for 5 min. The cells were then resuspended in 100 μL 20 mg/mL lysozyme solution and incubated for 20 min at room temperature. The cells were further lysed by 3 x 30 s bead-beating in the lysis buffer in RNeasy Mini Kit (Qiagen). Subsequent RNA extraction and purification was performed using the same kit. Total RNA was eluted from the spin column using nuclease-free water and stored at -80°C for further applications.

**RNA sequencing**

Total RNA samples with a minimum integrity number of 6 were used for downstream processes. The rRNA-depleting library preparation and sequencing were done by the integrated Genome Analytics Platform at Genome Institute of Singapore (Singapore). The library was sequenced on a HiSeq 4000 sequencing system (Illumina) with a pair-ended read length of 2*151 bp. Data analyses were done on the Galaxy server (https://usegalaxy.org.au/). Briefly, HISAT2 (4) was used to align the reads to the reference genome (GenBank NC_008596.1) and featureCounts (5) was used to measure the expression of each gene. Differential expression analysis was done using EdgeR (6).

**Quantitative real-time PCR**

500 ng isolated total RNA was converted to cDNA by reverse transcription using ReverTra Ace qPCR RT Master Mix with gDNA Remover (Toyobo). KOD SYBR qPCR Mix (Toyobo) was used for the PCR on an Applied Biosystems 7500 real-time PCR system (ThermoFisher Scientific). ROX was used as the reference dye and SYBR Green I included in the PCR Mix was used for the quantification. The qPCR

volume was 20 μL for each well with each primer at 200 nM. The transcript levels of *fas*, *inhA*, *kasA*, and *pks13* were quantified after normalization to the transcript level of housekeeping gene *sigA*. The thermocycling program for the PCR was as follows: 98 °C for 2 min followed by 40 cycles of 95 °C 20 sec, 60 °C 30 sec, and 68 °C 30 sec.

**SDS-PAGE of radiolabeled whole cell lysates**

1-mL stationary-phase TKO depleted of trehalose (dTKO) cells were radiolabeled with 2 μCi/mL [1-$^{14}$C]-acetate for 30 min and harvested by centrifugation at 4,000 x *g* for 3 min. 50 μL 2x non-reducing Laemmli buffer was added to the cell pellet. The sample was sonicated in a bath sonicator for 5 min and vortexed for 1 min after being mixed with glass beads. The sample was then heated at 95 °C for 10 min and centrifuged at 21,100 x *g* for 10 min. 20 μL sample was subjected to SDS-PAGE analysis using a 6% gel. After the electrophoresis, the gel was stained with InstantBlue Coomassie Protein Stain (Abcam) and dried using the DryEase Minigel Drying System (Invitrogen). The dried gel was then exposed to a storage phosphor screen for 3 days followed by visualization by phosphor imaging.

**In vitro mycolate release and reduction**

The *pks13* gene (*MSMEG_6392*) was cloned without the stop codon into pET22/42 digested with NdeI and HindIII via Gibson assembly. The gene was subcloned from there together with the upstream RBS and the downstream sequence encoding the C-terminal His-tag in pJEB402 digested with EcoRI and HindIII via Gibson assembly. TKO cells harboring a pJEB402-*pks13*-*his8* plasmid were grown from a single colony in 4 mL 7H9 medium in the presence of 5 μM trehalose. At the stationary phase, the culture was inoculated into 200 mL fresh medium containing 1 μM trehalose. The dTKO cells depleted of trehalose were harvested after 48 h by centrifugation at 4,000 x *g* for 2 min. Pks13-His$_8$ was purified by TALON affinity purification similar to that for CmrA-His$_8$ (see below). The protein was eluted from the affinity resin using TBS pH 7.4 containing 200 mM imidazole. The eluate was concentrated and dialyzed with TBS pH 7.4 using an Amicon Ultra 100 kDa centrifugal filter (MilliporeSigma) to remove excessive imidazole.

For the release assay, 5 mM of each putative acceptor molecules was added to purified Pks13-His$_8$. For reduction, 100 μM NADPH, 5 μg CmrA was added together with trehalose to a total 50 μL reaction mixture. The release/reduction mixture was kept at 37 °C for 30 min. To extract lipids from the solution, 300 μL methanol and 150 μL chloroform were added. After 10 s vortexing, 150 μL chloroform and 160 μL ultrapure water were added to allow phase separation. The bottom organic phase was collected before 300 μL chloroform was added to extract the lipids again. The two extracts were pooled and dried in a fumehood overnight. The dried lipids were reconstituted in 10 μL chloroform/methanol (2:1, v/v) and the entire sample (~1,000 cpm) was analyzed by TLC using a chloroform:methanol:water (30:8:1, v/v/v) solvent system. The dried TLC was then exposed to a storage phosphor screen for 5 days followed by visualization by phosphor imaging.

**Purification of CmrA**

The *cmrA* gene (*MSMEG_4722*) was cloned without the stop codon into pET22/42 digested with NdeI and XhoI via Gibson assembly. *E. coli* BL21(DE3) cells harboring the pET22/42-*cmrA*-*his$_8$* plasmid were grown from a single colony in 5 mL LB medium overnight. The culture was inoculated into 750 mL fresh medium and grown to $OD_{600}$ around 0.6. At this point, the protein expression was induced by 1 mM IPTG (isopropyl β-D-1-thiogalactopyranoside). The cells were harvested after 2 h by centrifugation at 4,800 x *g* for 10 min. The cells were resuspended in 10 mL cold TBS (Tris-buffered saline, 20 mM Tris-HCl pH 8.0, 150 mM NaCl) containing 100 μg/mL lysozyme (Calbiochem), 100 μM phenylmethylsulfonyl fluoride (PMSF; Calbiochem) and 50 μg/mL DNase I (MilliporeSigma), and lysed by two passages through a French Press (GlenMills) at 15,000 psi. The resulting suspension was centrifuged at 4,800 x *g* for 3 min to remove unbroken cells. The membrane debris was further removed by centrifugation at 100,000 x *g* for 30 min (Optima XL-90 ultracentrifuge, Beckman Coulter). The supernatant was loaded onto TALON metal affinity resin (Takara Bio) pre-equilibrated with TBS pH 8.0 containing 5 mM imidazole. The mixture was allowed to drain by gravity and the filtrate was loaded back to the column and drained again. The resin was washed 10 times with the washing buffer (20 mM Tris-HCl pH 8.0, 300 mM NaCl and 5 mM imidazole) and subsequently eluted with the elution buffer (TBS pH 8.0 containing 200 mM imidazole). The eluate was further purified by SEC (AKTA, GE Healthcare) at 4 °C on a pre-packed Superdex 75 10/300 GL column using TBS pH 8.0 as the eluent. Fractions corresponding to the peaks of our interest were concentrated using an Amicon Ultra 10 kDa centrifugal filter (MilliporeSigma), verified by SDS-PAGE analysis and stored in -80 °C for future usage.

**Drug sensitivity test**

Minimum inhibitory concentrations (MICs) of different drugs were measured in 7H9 liquid media in 96-well plates. TKO cells were grown with 1 μM trehalose to the stationary phase before dilution to $OD_{600}$ = 0.01 using fresh medium containing 10 μM trehalose or 6-azido-trehalose and aliquoted into the wells (100 μL each). Drugs of different concentrations were added into the wells following two-fold serial dilutions. The plate was incubated in a shaker at 37°C, 150 rpm for 36 h. Thiazolyl blue tetrazolium bromide (MTT; MilliporeSigma) was added to a final concentration of 0.2 mg/mL as the indicator of live cells. MIC was the minimal concentration of a drug on the plate that precluded any visible growth of the cells.

**Table S1.** MIC (μg/mL) of different drugs on different *M. smegmatis* strains.

| drug | WT | TKO (+tre)[a] | TKO (+Az)[a] |
|---|---|---|---|
| isoniazid | 2.5 | 2.5 | 2.5 |
| rifampicin | 1.25 | 0.625 | 0.625 |
| streptomycin | 0.25 | 0.25 | 0.125 |
| erythromycin | 10 | 1.25 | 2.5 |
| novobiocin | 4 | 1 | 1 |
| SDS[b] | 0.0125 | 0.0125 | 0.00625 |
| chloramphenicol | 30 | 30 | 30 |

[a] tre, trehalose; Az, 6-azido-trehalose.
[b] (%) instead of (μg/mL)

### Supplementary figures

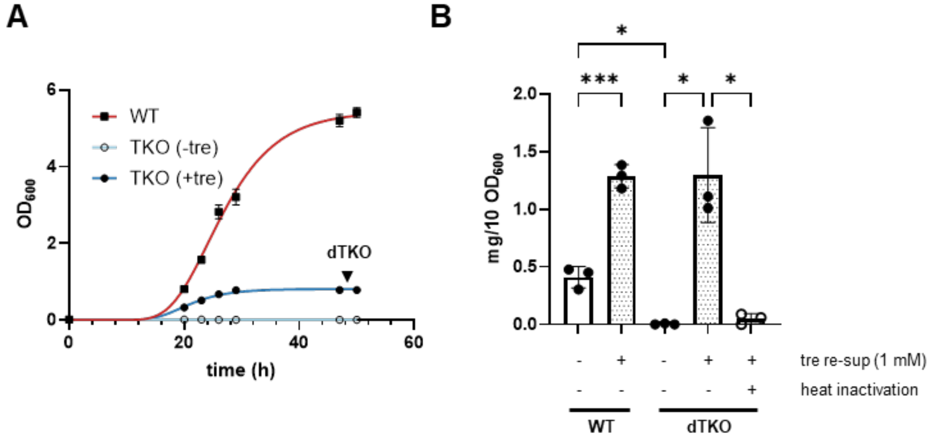


**Figure S1.** TKO cells stop growing when depleted of intracellular trehalose. (*A*) Growth curves of WT cells and TKO cells with or without trehalose. The TKO cells were depleted of trehalose before subculture into fresh medium with (+tre) or without (-tre) 1 µM trehalose. Each data point (mean ± SD) represents the results from three biological replicates. The black arrowhead indicates the time point when trehalose-depleted TKO (dTKO) cells were harvested and used for different experiments. (*B*) GC-MS quantification of intracellular trehalose of indicated strains with (+) or without (-) trehalose (1 mM) re-supplementation (tre re-sup). Each data point (mean ± SD) represents the results from three biological replicates. *, $p < 0.05$; ***, $p < 0.001$.

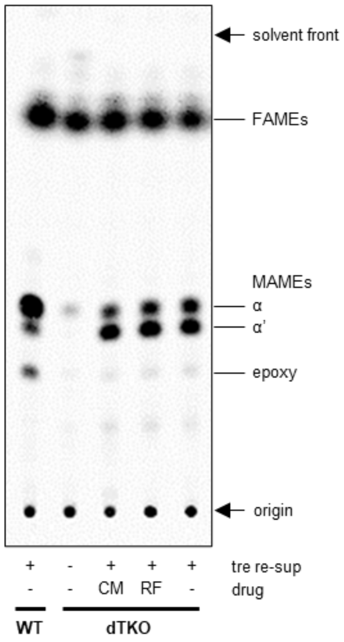


**Figure S2.** Trehalose re-supplementation restores MA biosynthesis in dTKO cells even when transcription or translation, thus cell growth, are inhibited. TLC analysis of [$^{14}$C]-labeled total acyl methyl esters extracted from *M. smegmatis* WT or dTKO (TKO depleted of trehalose) cells, with (+) or without (-) trehalose re-supplementation (tre re-sup). Chloramphenicol (CM) or rifampicin (RF) were added at 2x MIC concentrations to inhibit translation or transcription, respectively, during trehalose re-supplementation. The same amount of radioactivity was loaded for each sample. Developing solvent: three times in hexane:ethyl acetate 95:5. The TLC plate was visualized by phosphor imaging. FAME, fatty acid methyl ester; MAME, mycolic acid methyl ester.

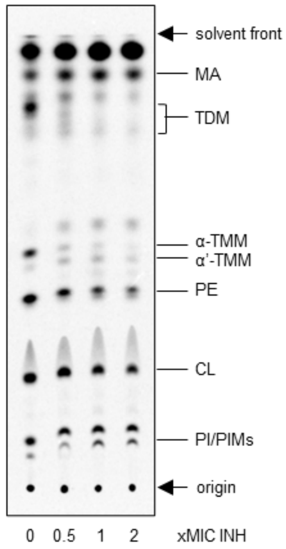


**Figure S3.** Treatment with FAS-II inhibitor isoniazid causes increase in the ratio of α'- to α-mycolates. TLC analysis of [$^{14}$C]-labeled lipids extracted from WT *M. smegmatis* cells treated with different concentrations of isoniazid (INH). Developing solvent: chloroform:methanol:water 30:8:1. The TLC plate was visualized by phosphor imaging. CL, cardiolipin; PE, phosphatidylethanolamine; PI, phosphatidylinositol; PIMs, phosphatidylinositol mannosides.

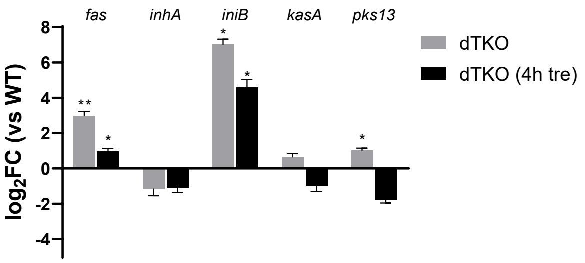


**Figure S4.** qRT-PCR validation of expression levels of indicated genes, when comparing dTKO cells (trehalose-depleted TKO), or dTKO cells with trehalose re-supplementation (4h tre), to WT cells.. The relative quantification was accomplished based on the $\Delta C_T$ (cycle threshold) after normalization to the housekeeping genes *sigA*. The values shown here represent the mean ± SD of three biological replicates. *, $p < 0.05$; **, $p < 0.01$ compared to WT cells.

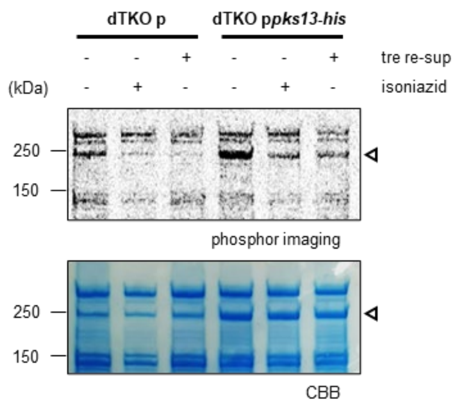


**Figure S5.** Trehalose re-supplementation or isoniazid treatment decreases [$^{14}C$]-acetate labeling of ~200 kDa protein band believed to be Pks13. SDS-PAGE analyses of lysate from dTKO cells harboring empty vector (dTKO p) or pJEB402-*pks13-his*$_8$ (dTKO p*pks13-his*) grown in the presence of [$^{14}C$]-acetate. The gel was visualized by phosphor imaging (top) and Coomassie brilliant blue staining (CBB; bottom). tre re-sup, trehalose re-supplementation. The results are representative of three biological replicates.

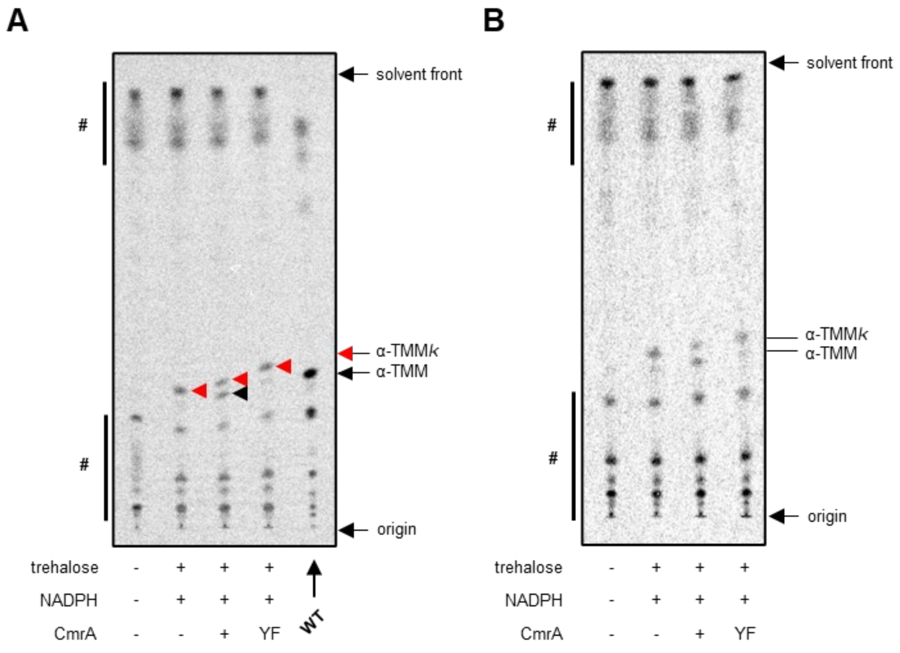


**Figure S6.** Replicate TLC analyses of lipids extracted from affinity-purified Pks13-His subjected to in vitro mycolate release by trehalose and reduction by CmrA, in addition to the TLC shown in Fig. 4*E*. Collectively, these biological triplicates (including TLC in Fig. 4*E*) were quantified, and the resulting data shown in Fig. 4*F*. The top and bottom spots (#) on the TLC other than TMM*k* and TMM are present even in the absence of trehalose and are likely non-specific lipids bound to TALON resin used for affinity purification of the protein. Developing solvent: chloroform:methanol:water 30:8:1. The TLC plates were visualized by phosphor imaging. YF, CmrA containing a Y157F mutation.

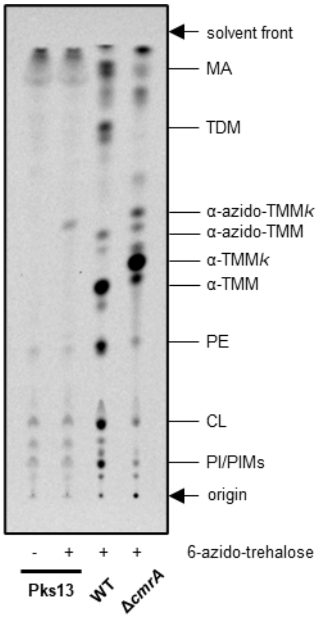


**Figure S7.** 6-azido-trehalose releases a lipid product from Pks13-His in vitro with similar migration as 6-azido-TMM*k*, produced in Δ*cmrA* cells fed with 6-azido-trehalose. TLC analysis of [$^{14}$C]-labeled lipids extracted from affinity-purified Pks13-His subjected to in vitro mycolate release by 6-azido-trehalose. Lipids extracted from WT and Δ*cmrA* cells fed with 6-azido-trehalose were used as references for migration positions of α-6-azido-TMM and α-6-azido-TMM*k*, respectively. For clarity, α'-(6-azido-)TMM and α'-(6-azido-)TMM*k* (spots immediately below corresponding α-(6-azido-)TMM and α-(6-azido-)TMM*k*) are not labelled for these samples. Developing solvent: chloroform:methanol:water 30:8:1. The TLC plate was visualized by phosphor imaging. CL, cardiolipin; PE, phosphatidylethanolamine; PI, phosphatidylinositol; PIMs, phosphatidylinositol mannosides.

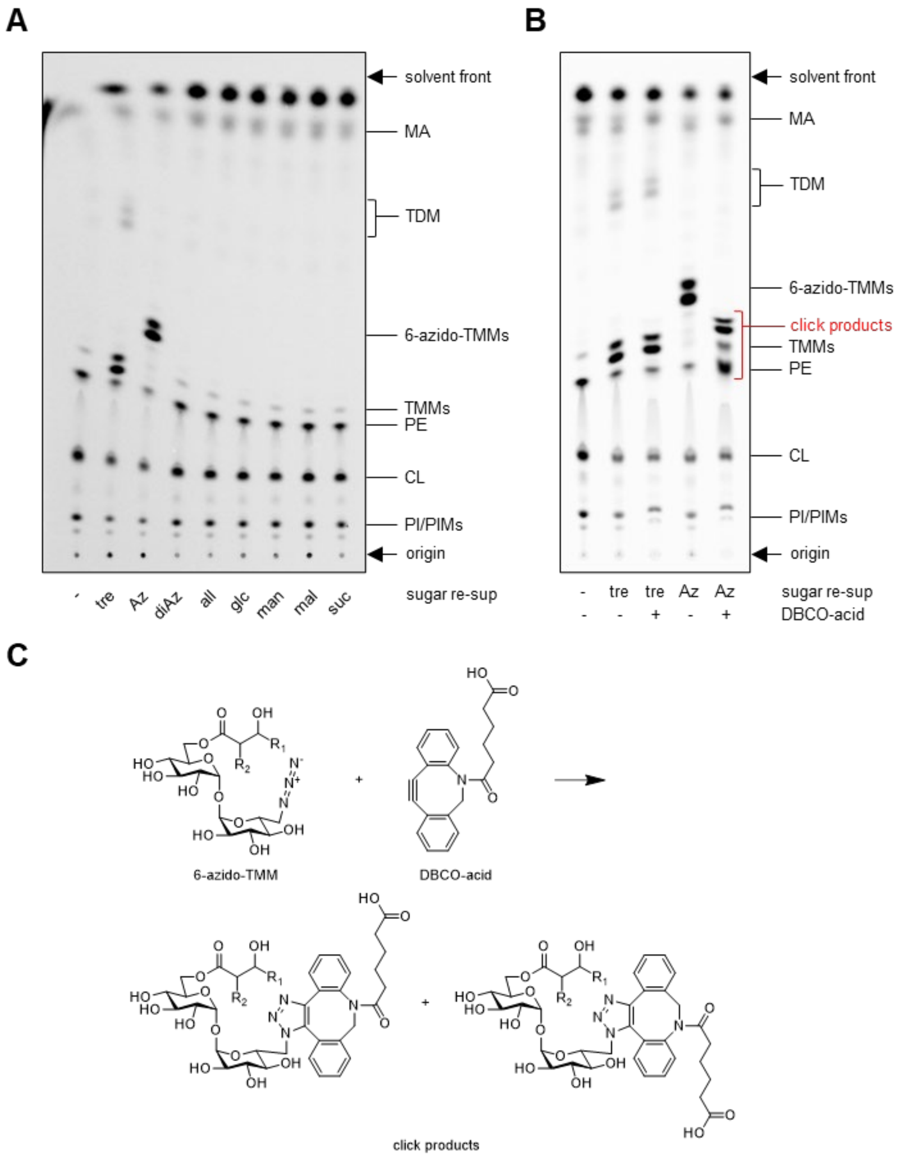


**Figure S8.** 6-azido-trehalose, but not other sugars, enabled the formation of newly-synthesized 6-azido-TMMs in trehalose-depleted TKO cells. (*A*) TLC analysis of [$^{14}$C]-labeled lipids extracted from dTKO cells re-supplemented with the indicated sugar. tre, trehalose; Az, 6-azido-trehalose; diAz, 6,6'-diazido-trehalose; all, allose; glc, glucose; man, mannose; mal, maltose; suc, sucrose. (*B*) TLC analysis of [$^{14}$C]-labeled lipids extracted from dTKO cells and reacted with DBCO-acid. Developing solvent: chloroform:methanol:water 30:8:1. The TLC plates were visualized by phosphor imaging. CL, cardiolipin; PE, phosphatidylethanolamine; PI, phosphatidylinositol; PIMs, phosphatidylinositol mannosides. (*C*) Reaction scheme illustrating strain-promoted [3+2] azide-alkyne cycloaddition ("click reaction") between 6-azido-

TMM and DBCO-acid.
